# Hybridization and cryptic speciation in the Tropical Eastern Pacific octocoral genus *Pacifigorgia*

**DOI:** 10.1101/2021.04.29.442007

**Authors:** Angelo Poliseno, Odalisca Breedy, Hector M. Guzman, Sergio Vargas

**Affiliations:** Department of Earth and Environmental Sciences, Geobiology & Paleontology, Ludwig-Maximilians-Universität München. Richard-Wagner-Str. 10, 80333 Munich, Germany; Centro de Investigación en Estructuras Microscópicas, Escuela de Biología, Universidad de Costa Rica. P.O. Box 11501-2060; Centro de Investigación en Ciencias del Mar y Limnología, Escuela de Biología, Universidad de Costa Rica. P.O. Box 11501-2060; Smithsonian Tropical Research Institute, P.O. Box 0843-03092, Panama, Republic of Panama

## Abstract

The shallow waters of the Tropical Eastern Pacific (TEP) harbor a species-rich octocoral fauna, with seven genera and 124 octocoral species described to date for the region. Of these lineages, *Pacifigorgia*, with 35 species, is by far the most speciose and abundant shallow-water octocoral occurring in the region. The speciation mechanisms resulting in this remarkable diversity remain speculative, despite the extensive taxonomic and molecular systematic research conducted so far in the TEP. Using genome-wide SNP markers, we provide evidence for hybridization and extensive cryptic speciation in *Pacifigorgia*, suggesting that the genus’ diversity has been underestimated by traditional and molecular systematic research. Our study highlights the difficulties faced by both traditional taxonomy and single-marker based molecular approaches to characterize octocoral diversity and evolution, and the role genome-wide molecular studies coupled to morphological research play to advance our understanding of this group.

## Introduction

Among Tropical Eastern Pacific (TEP) octocorals, *Pacifigorgia* (Fig. 1) represents the most species-rich genus with about 35 valid species described to date (Breedy and Guzman, 2003, 2002; Guzman and Breedy, 2012). About two-thirds *Pacifigorgia* species, including at least six endemic species restricted to the shallow waters of the Gulf of Chiriquí, Panama, occur in the Panamic province (from ∼16° N to ∼3° N) (Guzman et al., 2004; Vargas et al., 2008). This marked increase in species number toward lower latitudes in *Pacifigorgia* is consistent with the latitudinal diversity gradient reported for shallow-water eastern Pacific octocorals (Núñez–Flores et al., 2019) and likely drives it. Thus, clarifying the mechanisms of speciation that resulted in the high diversity of *Pacifigorgia* in the Panamic province is pivotal to understand how evolutionary processes shaped the octocoral diversity patterns observed in the region.

**Fig. 1:**
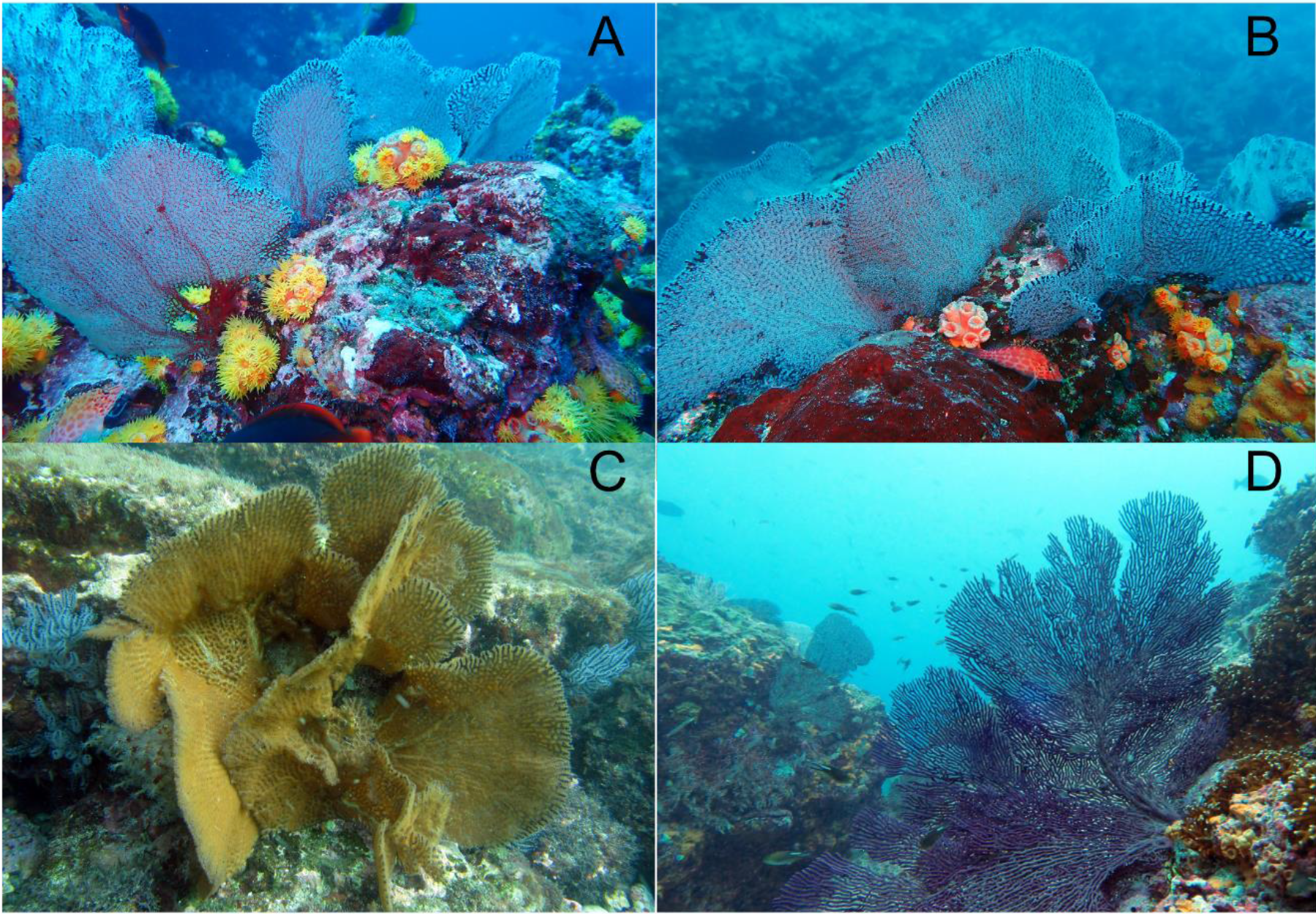
In situ photographs of A) *Pacifigorgia cairnsi*, B) *Pacifigorgia rubicunda*, C) *Pacifigorgia firma*, and D) *Pacifigorgia stenobrochis*. Photo credits: *P. cairnsi* and *P. rubicunda* Kike Ballesteros, *P. firma* Jaime Nivia, *P. stenobrochis* Kevan Mantell.

In contrast to most octocoral genera, *Pacifigorgia* has been the subject of intense morphological and molecular research. Breedy and Guzman (2002) thoroughly revised the genus and later described many new species (Breedy, 2001; Breedy and Guzman, 2004, 2003; Breedy and Guzmán, 2003; Guzman and Breedy, 2012). Molecular phylogenetic studies of *Pacifigorgia* are also available, yet species-level relationships within *Pacifigorgia* remain poorly resolved due to the lack of resolution of both mitochondrial (i.e., mtMutS) and nuclear markers (e.g., 28S rDNA) at this level (Ament-Velásquez et al., 2016; Soler-Hurtado et al., 2017; Vargas et al., 2014).

Despite the difficulties faced studying the diversification process in *Pacifigorgia* and other eastern Pacific octocorals (e.g., *Leptogorgia* and *Eugorgia*), some patterns arise from the molecular phylogenies available for the group. For instance, Ament et al. (2016) and Soler-Hurtado et al. (2017) proposed that hybridization could explain the mito-nuclear conflicts found in several eastern Pacific octocoral (holaxonian) genera. However, those authors’ inability to exclude other processes resulting in similar branching patterns, such as incomplete lineage sorting after rapid diversification events, left those claims mostly speculative. Similarly, hypotheses on cryptic speciation within *Pacifigorgia* are most likely affected by the resolution and the number of phylogenetic markers used, and by the differences in taxon-sampling across phylogenies inferred using different markers (Ament-Velásquez et al., 2016; Soler-Hurtado et al., 2017; Vargas et al., 2014). Thus, the contribution of these processes to the diversification of eastern Pacific octocorals remains to be determined.

Here, we use genome-wide, Single Nucleotide Polymorphisms (SNPs) and a collection of widespread and locally restricted *Pacifigorgia* species from the Gulf of Chiriquí, Panama, a biodiversity hot-spot for this genus (Guzman et al., 2004; Guzman and Breedy, 2008), to assess the contribution of hybridization and cryptic speciation to the diversification process in *Pacifigorgia*. We detected hybridization events and several instances of cryptic speciation among the *Pacifigorgia* species sampled. Our results provide conclusive evidence for reticulation among eastern Pacific octocorals and pose new challenges for better studying the diversity and distribution of these organisms.

## Materials and Methods

We collected by SCUBA diving 82 specimens belonging two genera and ten species (*P. bayeri, P. cairnsi, P. eximia, P. ferruginea, P. firma, P. rubicunda, P. smithsoniana, P. stenobrochis*, and *Leptogorgia pumila* and *Leptogorgia taboguillae*) from six different localities in the Coiba National Park, Panama (S. Fig. 1). Sampling depths ranged from 8 m to 24 m. Upon sampling, we sorted and morphologically identified all specimens in the field before preserving them in absolute ethanol until further processing.

We extracted gDNA using a standard CTAB protocol (Porebski et al., 1997), and quality controlled the extracts on 1.5% agarose gels. We checked the yield and purity of the extracts using a NanoDrop 2100. If needed, we digested RNA with RNase A and cleaned the resulting RNA-free extracts using a standard sodium acetate-ethanol precipitation. Of the 82 specimens extracted, only 40 yielded high molecular weight DNA. We used these specimens to prepare reduced representation libraries following the Genotyping-by-Sequencing (GBS) protocol of Elshire et al. (2011). Briefly, for each specimen, we digested ∼150 ng of gDNA with ApekI for two hours at 75°C and ligated the resulting fragments (one hour at 22°C) to one “common” and one barcoded adapter. We stopped the ligation reaction by heating the samples at 65°C for 30 minutes and we amplified (15 cycles; annealing temperature of 65°C and extension time of 30s) the adapter-ligated fragments using a universal non-barcoded primer (GBS_PrimerA) and a different barcoded primer for each sample. We purified the PCR products using 1.1 volume Agencourt AMPure XP beads (Beckman Coulter, Inc.) and quantified them using a QUBit® 2.0 fluorometer with a dsDNA HS Assay Kit (Invitrogen, Carlsbad, CA). Before sequencing, we pooled the libraries at equimolar concentrations and performed a final quality check using a Bioanalyzer 2100 (Agilent, Santa Clara, CA). The library pool was adjusted at a concentration of 10nM and sequenced on two lanes of an Illumina HiSeq 2500 (Illumina, San Diego, CA) using 100-bp single-end chemistry. The raw sequence reads are available under study accession PRJEB44220. We demultiplexed, quality controlled, filtered, and trimmed to 80-bp the ∼300×10^6^ sequence reads obtained from the two HiSeq lanes. We determined SNPs *de novo* with IPyRAD (Eaton and Overcast, 2020) using a clustering threshold of 0.85, maximum 20 SNPs and eight gaps per loci, and a minimum depth of six reads for base calling. These parameters have been successfully used in previous studies using RAD-Seq on octocorals (Herrera and Shank, 2016; e.g., Pante et al., 2015; Quattrini et al., 2019). For phylogenetic inference, we first produced an alignment containing loci present in at least 10% of the taxa (i.e., four specimens) and then discarded columns with >20% gaps to produce an alignment containing 122,464 sites present in at least 32 of 40 specimens. We used this alignment to infer a maximum likelihood phylogeny in the program RAxML v8.2.12 (single partition GTRGAMMA+F, 1000 fast bootstrap replicates, Stamatakis, 2014). We estimated a SNP matrix using the same parameters described above but including only those SNPs present in at least 50% of the taxa (i.e., 20 specimens). We used this matrix to find specimen groups in a phylogenetic independent way using Discriminant Analysis of Principal Components (DAPC) analyses (Jombart et al., 2010). We used the package *adegenet* (Jombart, 2008), 10 Principal Components, and the method *find*.*clusters* to select the most probable number of species groups via BIC. For each specimen, we also calculated its group assignment probability. To test for hybridization in *Pacifigorgia*, we used IPyRAD to conduct ABBA-BABA tests on the best maximum likelihood phylogeny and the SNPs dataset estimated for phylogeny-independent analyses. The data matrices are publicly available at https://gitlab.lrz.de/palmuc/pacifigorgia-gbs

## Results and Discussion

Previous molecular phylogenetic studies of eastern Pacific octocorals recovered *Pacifigorgia* monophyletic but could not resolve its species-level relationships (Ament-Velásquez et al., 2016; Soler-Hurtado et al., 2017; Vargas et al., 2014). In contrast, we recovered a maximum likelihood phylogeny generally showing branch support values >75% (Fig. 2) and clades corresponding to recently revised and described *Pacifigorgia* species, such as *P. eximia* and *P. bayeri*. These species are morphologically well-defined and can be accurately identified in the field and laboratory (Breedy and Guzman, 2004, 2003). Accordingly, these species belonged to clearly defined DAPC groups, and we could not detect introgression between these species and specimens of other *Pacifigorgia* clades. Thus, our genome-wide GBS data further support their species-level status.

**Fig. 2:**
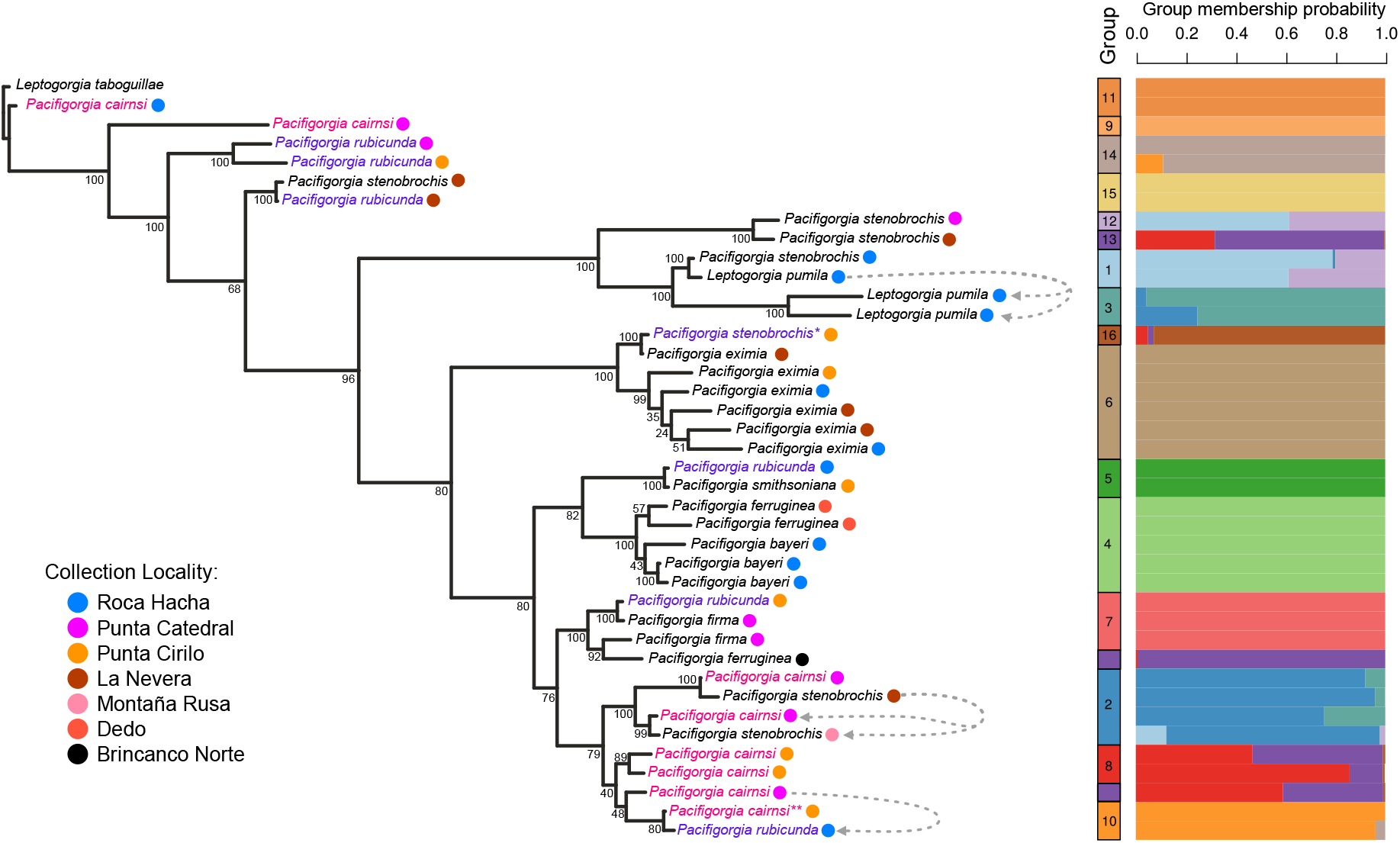
Maximum likelihood phylogeny of *Pacifigorgia* based on 122,464 genome-wide SNP markers. Numbers below the branches represent bootstrap support values. *Pacifigorgia cairnsi* and *Pacifigorgia rubicunda*, two widespread species, are highlighted in light red and purple blue, respectively. Gray dashed arrows indicate introgression events detected using ABBA-BABA tests on the maximum-likelihood topology. On the right, group assignments and group membership probabilities of different *Pacifigorgia* specimens obtained using DAPC of the SNPs matrix. \**P. stenobrochis* and \*\**P. cairnsi* colonies with anomalous sclerome and DAPC group assignments.

Our phylogeny also recovered well-supported clades grouping individuals assigned to different morphologically defined species. For instance, we detected a group composed of individuals assigned to *P. stenobrochis* and *Leptogorgia pumila* (Fig. 2). Ament et al. (2016) also found this same grouping using partial mtMutS and 28S rDNA sequences and proposed to transfer *L. pumila* to *Pacifigorgia*. However, such taxonomic decision requires the amendment of *Pacifigorgia*’s diagnosis to include non-anastomosing or loosely anastomosing species, an act that will let the genus so poorly defined that it could, in principle, include all *Leptogorgia* species hitherto described. Within this clade, DAPC assigned *L. pumila* to groups one and three and *P. stenobrochis* specimens to groups one, twelve, and thirteen (Fig. 2). Groups one and twelve likely are in admixture; *P. stenobrochis* specimens have high group membership probability for either group. Group thirteen appears to encompass specimens with high levels of missing data in the SNP matrix (S. Fig 2) and should be taken cautiously. We detected introgression between all specimens identified morphologically as *L. pumila* (Fig. 2 and S.Fig. 3) despite the phylogenetic tree pointing to a closer phylogenetic relationship between some *P. stenobrochis* and *L. pumila* specimens and the assignment of *L. pumila* specimens to different DAPC groups.

Similarly, we detected a well-supported clade grouping specimens of *P. carnsi* and *P. stenobrochis* (Fig. 2). DAPC assigned all four specimens included in this clade to the same group, and we detected introgression between them (S. Fig. 4). These results indicate that this group’s specimens, although assigned to different morphological species, are in admixture. Previous DNA barcoding studies also revealed multiple *P. stenobrochis* clades in the eastern Pacific (Vargas et al., 2014). We recovered four not closely-related clades and an equal number of DAPC groups including *P. stenobrochis* specimens, further pointing to the existence of cryptic lineages within this species. Some of these clades and groups include “typical” *P. stenobrochis* individuals while other consist of specimens differing in colony shape or spiculation from this species’ type (e.g. DAPC Group 16 *P. stenobrochis*). Despite its very characteristic colony morphology, *P. stenobrochis* displays some variation in the mesh, and two recognized color morphs can be observed in the same colony. The species’ sclerome is also variable, with either spindles or blunt spindles as the dominant sclerite form (Breedy and Guzman, 2004, 2002). However, in phenotypically very plastic organisms like octocorals (Calixto-Botía and Sánchez, 2017; Prada et al., 2008; West et al., 1993), it is hard to justify solely based on morphology the segregation of *P. stenobrochis* into multiple species with slightly different coloration, sclerome or colony shapes, or based on the occasional collection of specimens not fitting perfectly the *P. stenobrochis* gestalt. Our results indicate that a taxonomic reevaluation of *P. stenobrochis*’ different morphs is warranted and provide an integrative framework to morphologically describe new species leveraging high-resolution, genome-wide markers.

Our phylogenetic results supported clades including specimens of *P. rubicunda* and *P. cairnsi*, and of *P. rubicunda* and *P. stenobrochis* (Fig. 2). These clades also corresponded to DAPC groups (i.e., groups 10 and 15). The morphology of the specimens included in these groups was typical; the specimen of *P. cairnsi* had a somewhat divergent sclerome dominated by spindles. *Pacifigorgia rubicunda* is morphologically diverse, with colonies consisting of single fans or forming rosettes. This species coexists with *P. cairnsi* and *P. stenobrochis* throughout its geographic range (Breedy and Guzman, 2003, 2002). We found evidence for introgression between *P. cairnsi* and *P. rubicunda* (Fig. 2 and S. Fig. 5), which are morphologically different. However, this hybridization event involves one *P. cairnsi* specimen assigned to group thirteen, a group joining mainly specimens with many missing SNPs, and a clade with internal low bootstrap support values (Fig. 2). Therefore, this introgression event should be taken cautiously. Beside these mixed clades, we found highly supported clades of *P. cairnsi* and *P. rubicunda* specimens collected in different localities interspersed in the tree (Fig. 2). DAPC also supported these clades (e.g., group 14). We could not detect introgression for these clades, which likely represent cryptic lineages within *P. cairnsi* and *P. rubicunda*, two species with a wide geographic distribution (Breedy and Guzman, 2003).

We observed a similar phenomenon for the specimens of *P. firma* included in the analyses, which did not form a clade but grouped with specimens of *P. rubicunda* and *P. ferruginea* (Fig. 2). Except for *P. ferruginea*, DAPC assigned this clade’s specimens to group seven. We could not find any evidence for introgression within this clade, suggesting that *P. ferruginea* and *P. firma* currently include multiple evolutionary lineages (S. Fig. 6). *Pacifigorgia ferruginea* is endemic to the Gulf of Chiriqui and generally easy to identify in the field by its characteristic rusty appearance. In our phylogenetic analysis, *P. ferruginea* specimens from Dedo did not group with the single specimen collected in the nearby Brincanco Norte island, suggesting that cryptic species can occur within potentially very short geographic scales in *Pacifigorgia*. However, given the assignment of the single specimen of *P. ferruginea* from Brincanco Norte to DAPC group thirteen and a large number of missing SNP loci in this specimen, this interpretation should be corroborated in future studies. *Pacifigorgia firma* is widely distributed and morphologically variable (Breedy and Guzman, 2003). Despite its morphological plasticity and the continuous nature of most characters used to define the species, *P. firma* can be accurately determined. The finding of specimens morphologically assigned to *P. rubicunda* within a highly supported *P. firma* clade and DAPC highlights the challenges in establishing a morphological-molecular classification of *Pacifigorgia*. In particular, in morphologically highly variable and widespread species such as *P. rubicunda*, defining the extent to which hybridization contributes to the generation of new *Pacifigorgia* lineages and the molecular and morphological boundaries of those species remains to be determined.

Hybridization occurs in several scleractinian coral and octocoral lineages and is thought to be an important speciation mechanism in these groups (Mao, 2020). In scleractinian corals, hybridization between species with overlapping distribution ranges explains the large number of morphological species observed in important West and East Pacific reef-builders such as *Acropora* (Van Oppen et al., 2002) and *Pocillopora* (Combosch and Vollmer, 2015), respectively. In octocorals, McFadden and Hutchinson (2004) found evidence for hybridization in *Alcyonium* and Quattrini et al. (2019) found evidence of both morphological diversification in the absence of molecular divergence and hybridization in *Sinularia*. In this genus, local hybrids can become highly abundant, replacing their parental lines and changing community composition overtime (Slattery et al., 2008). Thus, other than evolutionary dead-ends, coral and octocoral hybrids represent evolutionary experiments capable of affecting the entire ecosystem. In the specific case of *Pacifigorgia*, the poor congruence between some morphological species and molecular groups we inferred using an extensive, genome-wide marker set indicates that genotyping specimens throughout a species’ geographic range is necessary to unveil morphologically cryptic and hybrid *Pacifigorgia* lineages and uncover the “true” diversity of this species-rich genus. Additionally, identifying differences in the reproductive cycles and strategies among different *Pacifigorgia* species and their populations is of crucial importance for better linking the molecular results with the reproductive ecology and natural history of these organisms (Gomez et al., 2018). In conjunction, our study suggests that the diversity of eastern Pacific *Pacifigorgia* is larger than currently recognized. Neutral processes such as the mid-domain effect cannot explain the diversity patterns observed for eastern Pacific octocorals (Núñez–Flores et al., 2019). Our data indicate that hybridization and cryptic speciation shape *Pacifigorgia*’s diversification history and likely are the drivers of the octocoral diversity patterns observed in the Tropical Eastern Pacific.

## Acknowledgments

We thank Andrea Quattrini for commenting an early version of the manuscript and providing valuable feedback on the interpretation of the analyses. We thank Prof. Gert Wörheide for providing access to laboratory facilities at the Dept. of Earth and Environmental Sciences, Geobiology & Paleontology (Ludwig-Maximilians-Universität München). We thank Dr. Stefan Krebs and Dr. Helmuth Blum (Ludwig-Maximilians-Universität, München) for technical support and sequencing the GBS libraries. The project was partially supported by the LMU München German Excellence Initiative Junior Research Funds to SV, the Smithsonian Tropical Research Institute, and the Secretaria Nacional de Ciencias y Tecnologia (SENACYT) de Panamá to HMG. The Ministerio de Ambiente de Panamá issued collection and exporting permits of eastern Pacific material. We thank C. Guevara and K. Mantell for their support during field collections. SV thanks N. Villalobos Trigueros, M. Vargas Villalobos, S. Vargas Villalobos, and S. Vargas Villalobos for their constant support.

## Author contributions

**AP:** Investigation, Formal Analysis, Data curation, Visualization, Writing – Original Draft. **OB:** Conceptualization, Investigation, Resources, Writing – Review & Editing, Project Administration. **HMG:** Resources, Funding Acquisition, Writing – Review & Editing, Project Administration. **SV:** Conceptualization, Methodology, Formal Analysis, Visualization, Writing – Original Draft, Resources, Supervision, Project Administration, Funding Acquisition.

## Notes

https://gitlab.lrz.de/palmuc/pacifigorgia-gbs

### Competing Interest Statement

The authors have declared no competing interest.

