## Supplementary Figures and Tables for "Hybridization and cryptic speciation in the Tropical Eastern Pacific octocoral genus *Pacifigorgia*"

**Supplementary Figure Legends**

Suppl. Fig. 1: Map of the Chiriquí Gulf showing the collection localities of the samples used in this study.

Suppl. Fig. 2: Matrix showing missing data (white) for *Pacifigorgia* SNPs. The arrows point specimens assigned to Group 13.

Suppl. Fig. 3: Results of the ABBA-BABA test for the *P. stenobrochis* + *L. pumila* clade.

Suppl. Fig. 4: Results of the ABBA-BABA test for the *P. stenobrochis* + *P. cairnsi* clade.

Suppl. Fig. 5: Results of the ABBA-BABA test for the *P. cairnsi* + *P. rubicunda* clade.

Suppl. Fig. 6: Results of the ABBA-BABA test for the *P. firma*+ *P. rubicunda* + *P. ferruginea* clade.

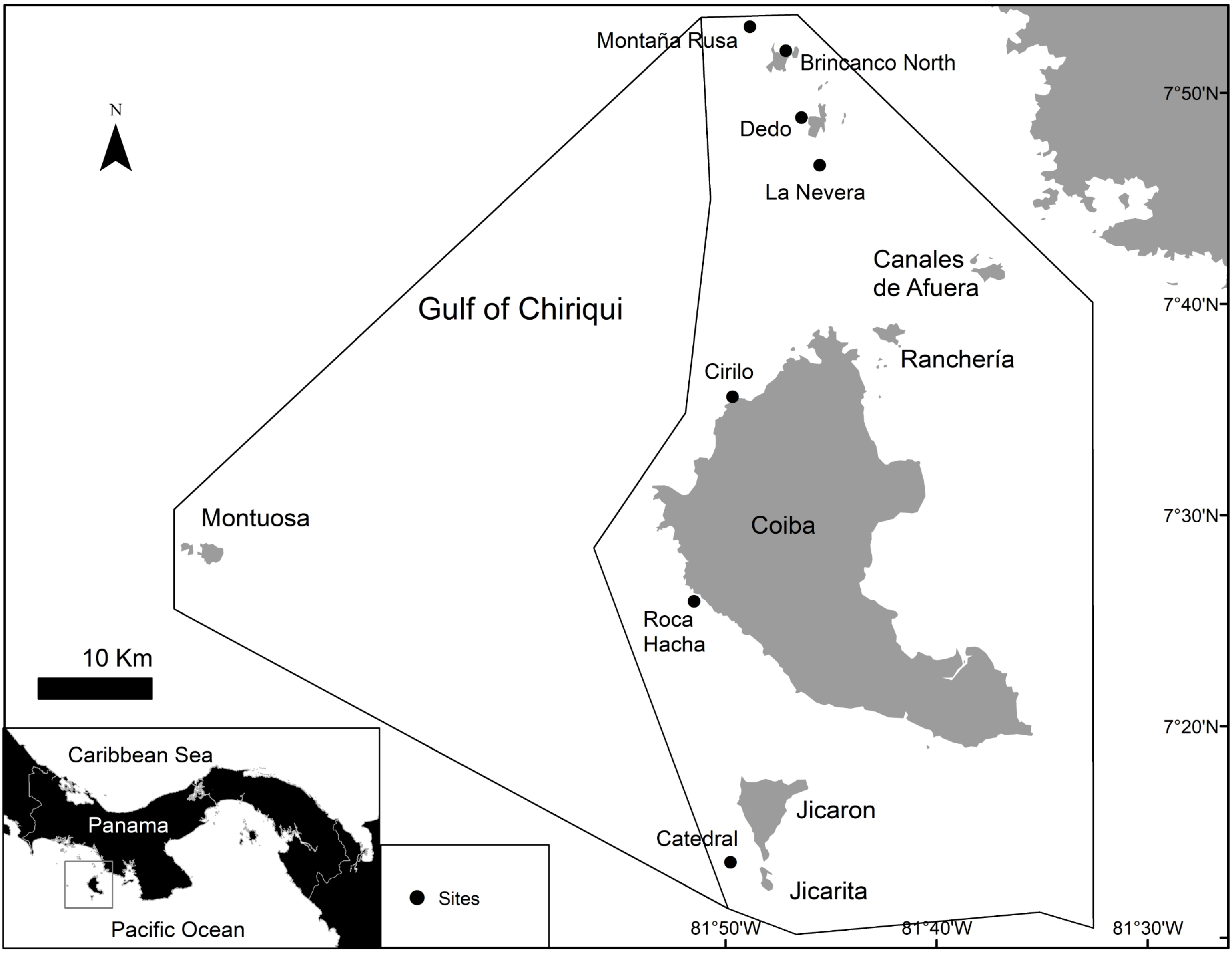

N
Montaña Rusa
Brincanco North
Dedo
La Nevera
Canales de Afuera
Gulf of Chiriqui
Ranchería
Cirilo
Montuosa
Coiba
Roca Hacha
10 Km
Caribbean Sea
Panama
Pacific Ocean
Jicaron
Catedral
Jicarita
Sites
7°50'N
7°40'N
7°30'N
7°20'N
81°50'W
81°40'W
81°30'W

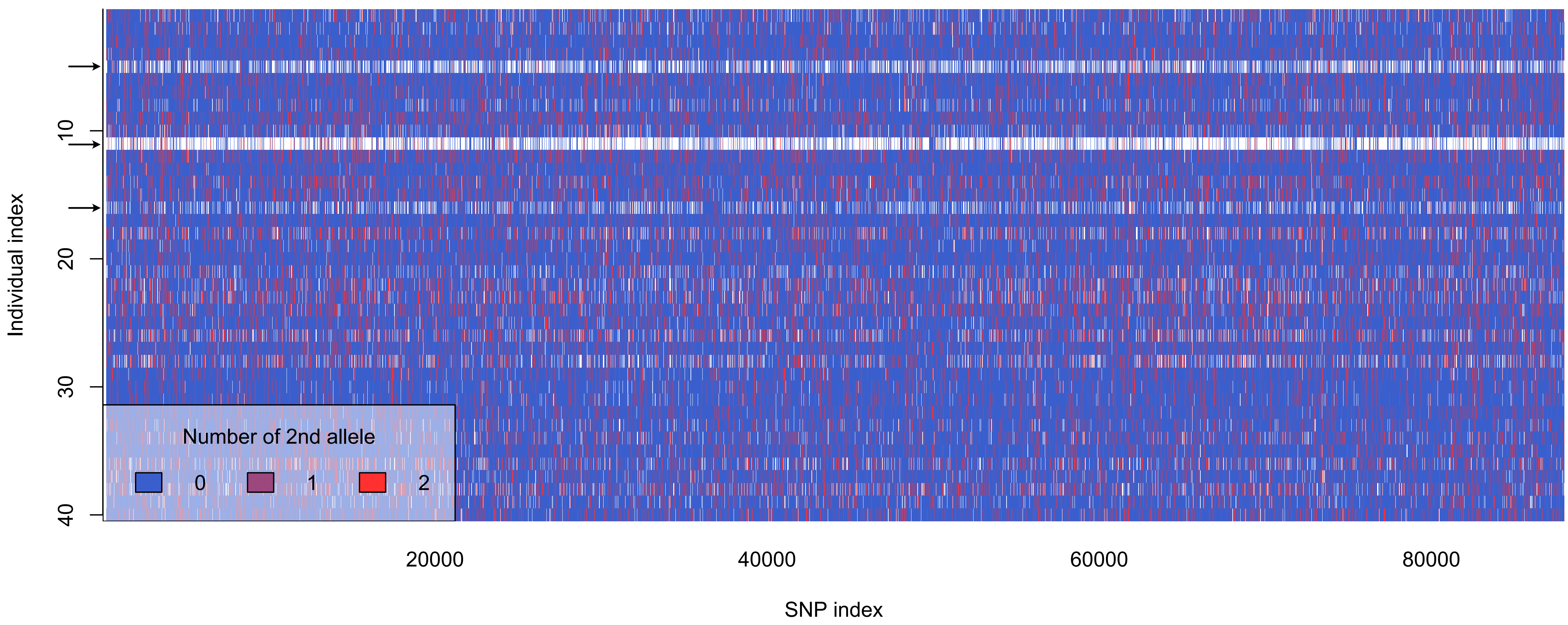

Number of 2nd allele
0
1
2
Individual index
10
20
30
40
SNP index
20000
40000
60000
80000

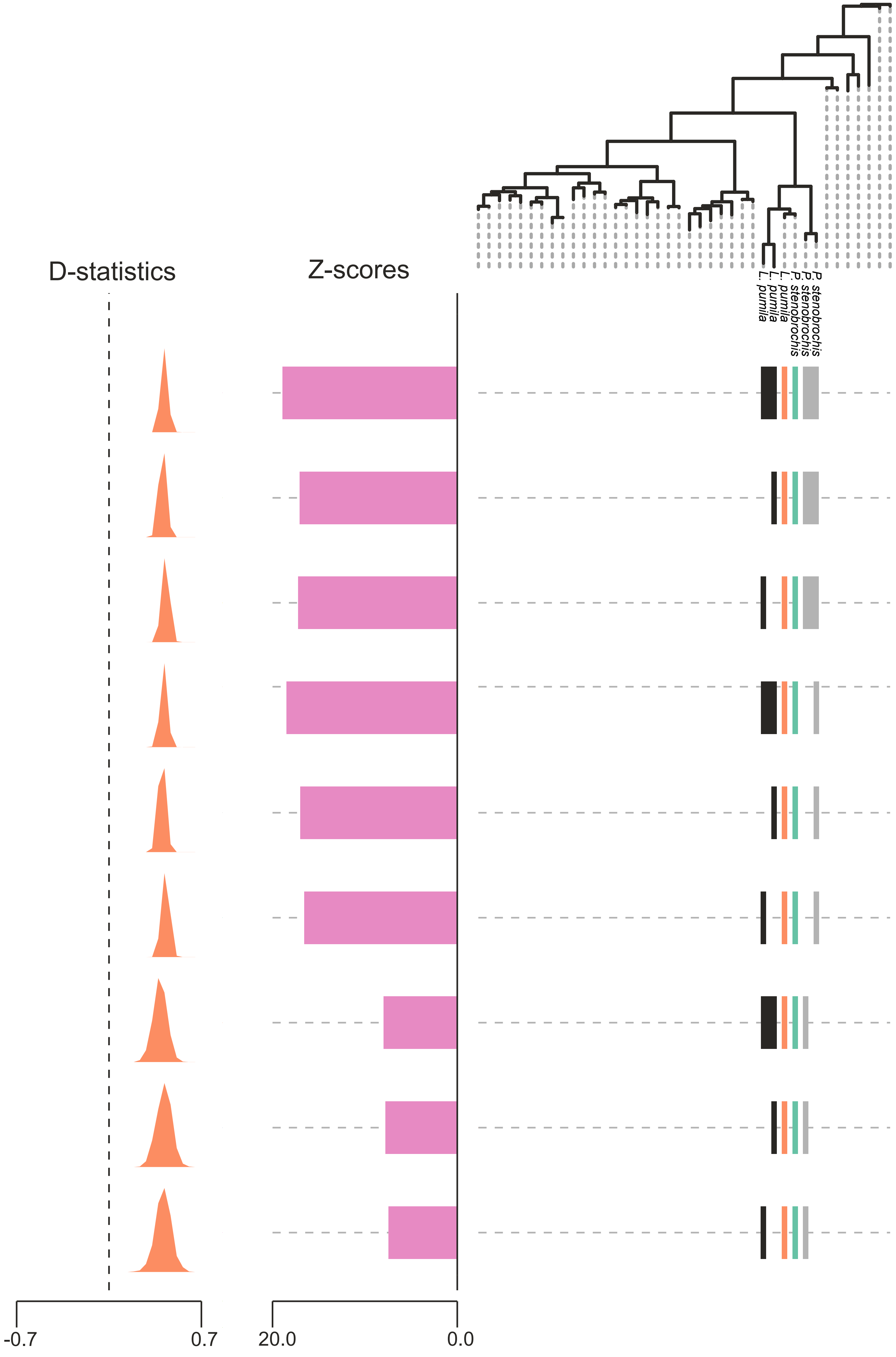

D-statistics
Z-scores
L. pumila
L. pumila
L. pumila
P. stenobrochis
P. stenobrochis
P. stenobrochis
-0.7
0.7
20.0
0.0

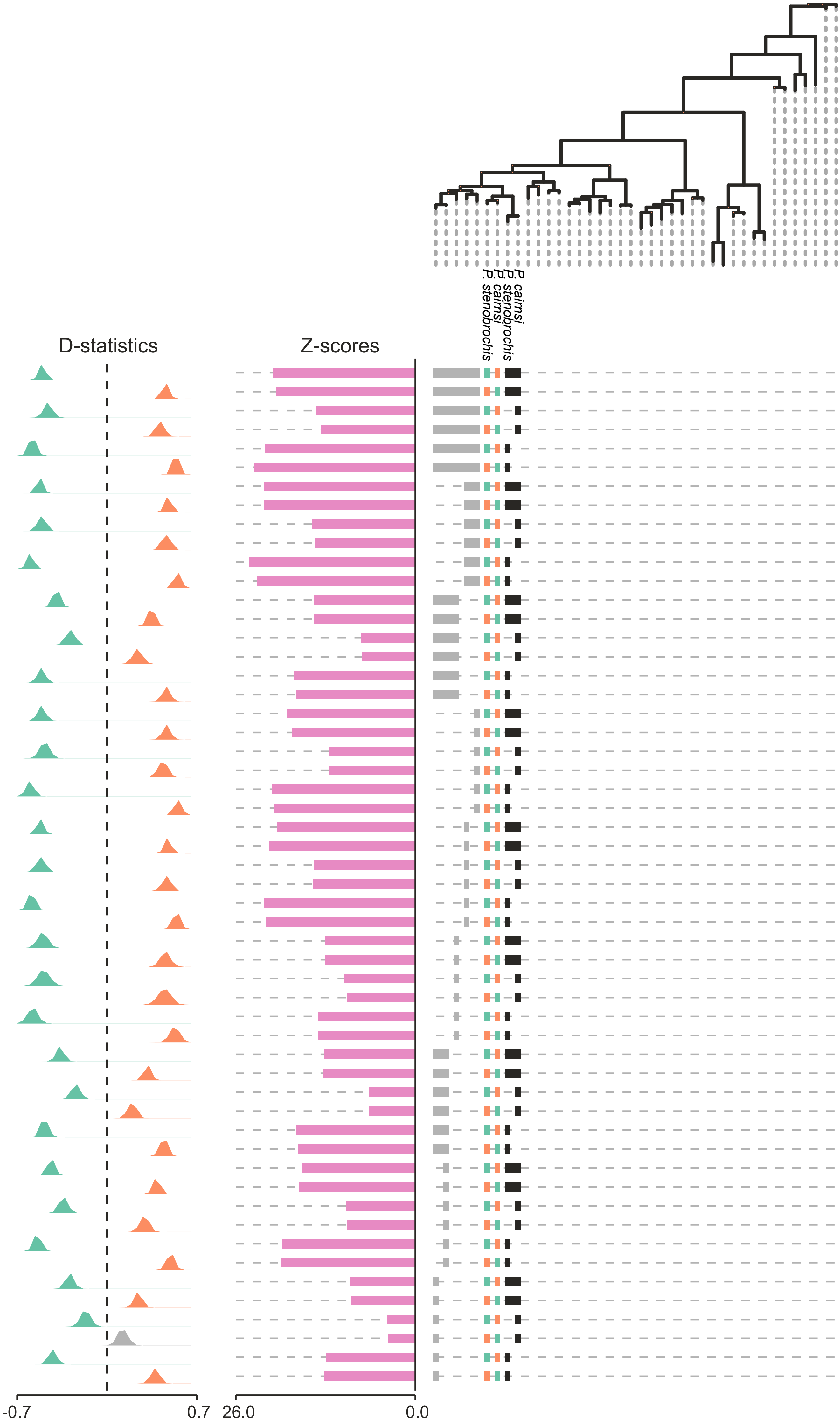

P. cairnsi
P. stenobrochis
P. cairnsi
P. stenobrochis
D-statistics
Z-scores
-0.7
0.7
26.0
0.0

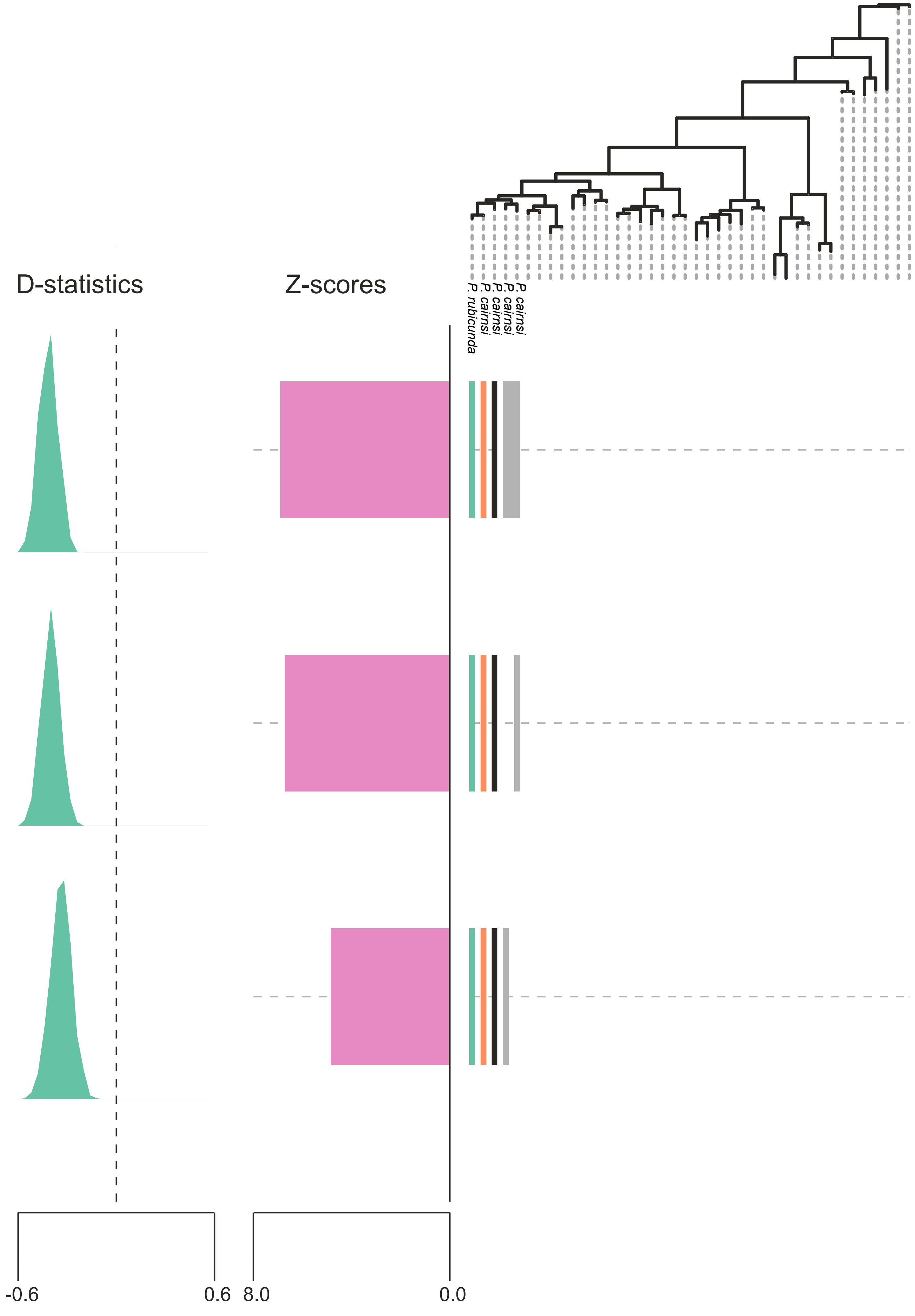
D-statistics
Z-scores
P. rubicunda
P. cairnsi
P. cairnsi
P. cairnsi
P. cairnsi
-0.6
0.6
8.0
0.0

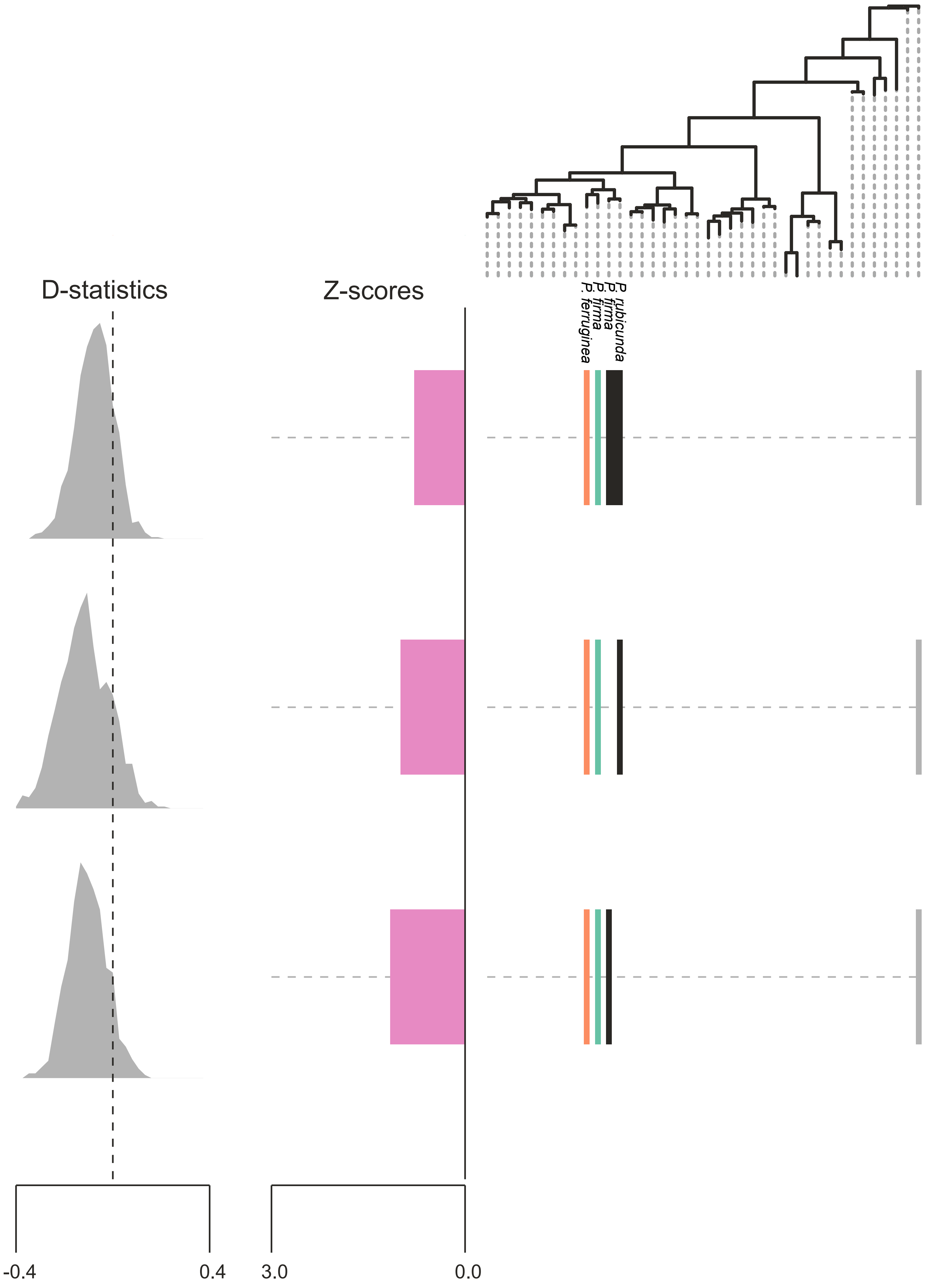
D-statistics
Z-scores
P. ferruginea
P. firma
P. firma
P. rubicunda
-0.4
0.4
3.0
0.0
